# Tumor fitness, immune exhaustion and clinical outcomes: impact of immune checkpoint inhibitors

**DOI:** 10.1101/679886

**Authors:** Adrian Bubie, Edgar Gonzalez-Kozlova, Nicholas Akers, Augusto Villanueva, Bojan Losic

## Abstract

Recently proposed tumor fitness measures, based on profiling neoepitopes for reactive viral epitope similarity, have been proposed to predict response to immune checkpoint inhibitors in melanoma and small-cell lung cancer. Here we apply these checkpoint based fitness measures measures to the matched checkpoint treatment naive TCGA samples where cytolytic activity imparts a known survival benefit. We observed no significant survival predictive power beyond that of overall patient tumor mutation burden, and furthermore, found no association between checkpoint based fitness and tumor T-cell infiltration, cytolytic activity (CYT), and abundance (TIL burden). In addition, we investigated the key assumption of viral epitope similarity driving immune response in the hepatitis B virally infected liver cancer TCGA cohort, and uncover suggestive evidence that tumor neoepitopes actually dominate viral epitopes in putative immunogenicity and plausibly drive immune response and recruitment.

## Introduction

Quantifying tumor fitness modulation by immune selection pressure has assumed a new urgency given the growing need to predict immunotherapy efficacy^1,2^. Immune-based anti-tumoral response is driven in part by T-cell receptor (TCR) recognition of tumor neoepitopes, which mainly represent mutated protein epitopes (i.e., neoepitopes) derived from somatic mutations. These neoepitopes are presented by host-cell major histocompatibility complex (MHC) molecules to facilitate ‘self’ recognition by TCRs (**Fig. 1a**). Cancer clones that escape immune recognition can hijack CTLA-4 and PD-1 checkpoint blockade, resulting in heterogeneous tumors with a complex immunogenomic profile which may include immune exhaustion, giving rise to mixed or unfavorable response to immune checkpoint inhibition^1,3^.

**Figure 1:**
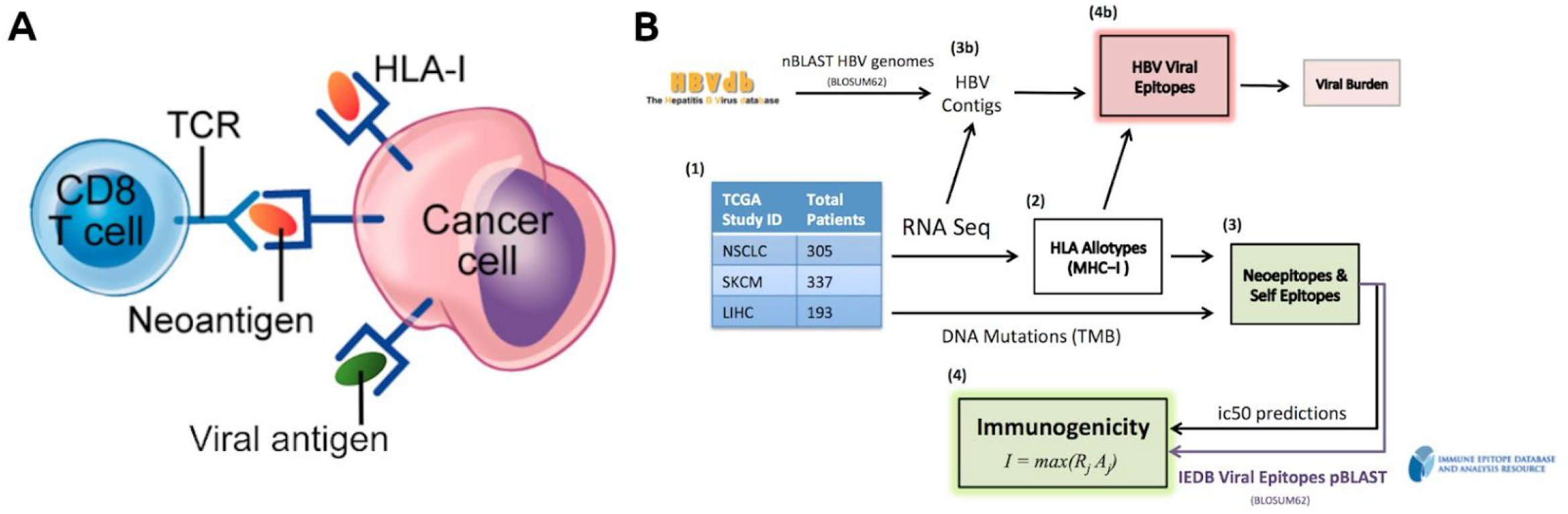
**A)** T-cell recognition by TCR of a cancer cell by MHC-I presentation. Antigens bound to MHC can displace other antigens due to higher binding affinity with the MHC molecule. This competitive binding affinity is represented by an ic50 score. **B)** Workflow summary. Matched mutation and RNA-seq TCGA datasets underwent HLA-Typing, neoepitope calling, and VDJ-alignment for each cohort, and tumor fitness scores calculated for those in the SKCM and NSCLC. Unmapped reads from the LIHC cohort were mined for HBV reads, which were used to predict viral epitopes. Logo provided by IEDB^5^.

Recently, a tumor fitness model has been proposed that incorporates two key components defining the TCR-neoepitope interaction: putative neoepitope immunogenicity and TCR recognition^4^. Putative immunogenicity is modeled via a nonlinear function of *in silico* derived MHC-I binding affinities arising from the clonal somatic mutation spectrum, while the TCR recognition likelihood is set to scale with sequence similarity to known, infectious, virally derived epitopes from the Immune Epitope Database (IEDB)^5^. Given these assumptions, a combined measure of tumor fitness is then defined as the inverse of the maximum clonal immunogenicity potential, Ϊ, weighted across subclones in the tumor (see Methods, **Fig. 1b**).

Although this model successfully predicted response to checkpoint inhibition in melanoma and small cell lung cancer, it is of interest to investigate its applicability, positive predictive power, and key assumptions in the arena of the analogous immunotherapy naive (endogenous) TCGA skin-cutaneous and melanoma (SKCM) and non-small lung cancer (NSCLC) cohorts. Crucially, even though checkpoint response predictive biomarkers are not necessarily survival prognostic, substantial evidence has already accumulated about the positive survival effects of cytolytic immune activity in melanoma and lung cancer^6,7^, strongly suggesting immune selection pressure significantly modulates treatment naive tumor fitness. Our central hypothesis was that the predictive tumor fitness model of checkpoint response will be survival prognostic in cohorts where cytolytic activity imparts a significant survival benefit. Indeed, the tumor fitness model makes no mathematical distinction between endogenous or checkpoint-induced immune response^4^.

An important corollary to our hypothesis was to directly test a key, central assumption of the *Luksza et al.* model, namely that viral similarity drives differential immune recognition of neoepitopes^4^. We hypothesized that in the setting of virally infected tumors, with the simultaneous presence of viral epitopes and neoepitopes, we could directly determine if the predicted MHC-I binding affinity (putative immunogenicity) of viral peptides was higher, lower, or indistinguishable from that of tumor neoepitopes *from the same patients*. Developing and deploying new in-silico methods, we carried this test out using the hepatitis B virally positive portion (HBV+) of the LIHC TCGA liver cancer cohort.

In this brief report we show that there is no significant correlation between tumor fitness and immune response signatures, nor is patient survival predicted using tumor fitness better than with ordinary tumor mutation burden (TMB), in either SKCM or NSCLC cohorts. Indeed we show that, as demonstrated in the checkpoint treatment setting^8^, a simple expression signature capturing tumor-infiltrating T-cell activity (CYT) significantly outperforms TMB in predicting survival in SKCM, and nearly so in NSCLC. We also show that tumor neoepitopes actually dominate viral epitopes in average binding affinity modulated immunogenicity within the same patients, and also better correlate with immune response, in the HBV+ HCC LIHC cohort.

## Results

### CYT, not Tumor fitness, correlates with immune measures in NSCLC and SKCM

We selected 305 patients with non-small cell lung cancer (NSCLC) and 337 with skin cutaneous and melanoma (SKCM) in the TCGA with matched whole exome sequencing (WES) and RNA-seq data (**Fig. 1b**). Putative tumor neoepitopes (ic50 < 500nM) with significant homology with viral sequences (e-value < 10) from the IEDB (see Methods) were found in 204/337 (60%) of SKCM and 221/306 (72%) of NSCLC patients. We found that SKCM patients with viral-like epitopes had a significantly worse survival rate, and correlated with lower TMB (but not CYT), while no survival associations were observed for NSCLC (**Supp. Fig. 3**). We observed significant associations between CYT and TIL burden for both the NSCLC and SKCM cohorts (**Fig. 2a,b**), as previously observed in checkpoint treated matched cohorts^8^. Significantly, however, we found that maximal clonal immunogenicity Ϊ does not correlate with TIL burden or CYT in either cohort.

**Figure 2:**
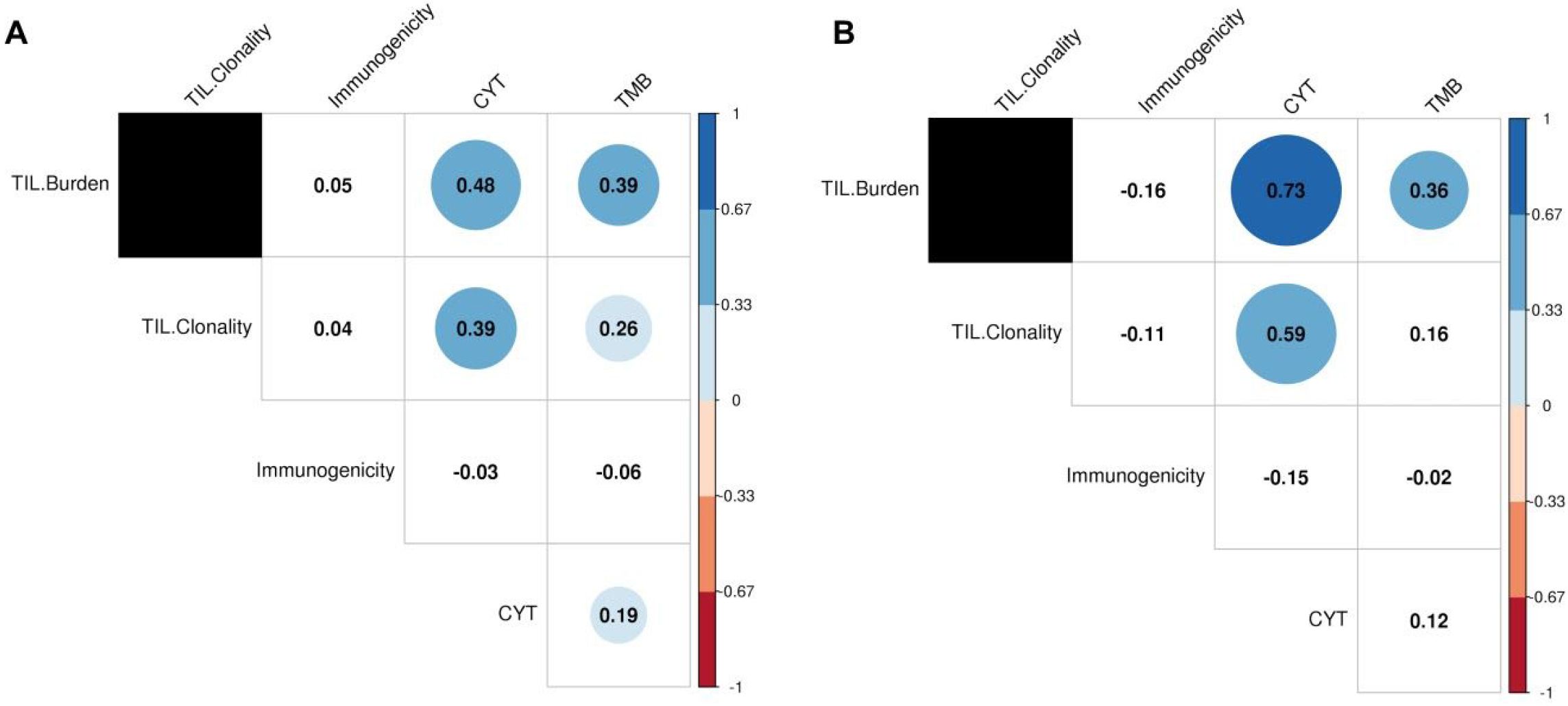
Maximal clonal immunogenicity (inverse tumor fitness), Ϊ, was found to be uncorrelated with measures of expression-based tumor immune response (TIL burden, TIL clonality, and CYT) and TMB in both **A)** NSCLC, **B)** SKCM patients. Several of the immune measures were found to be highly correlated with each other in both TCGA patient sets. Values in each circle indicate the Spearman correlations between the features, with color and size representative of correlation strength. Tiles with no color indicate a non-significant (p > 0.05) correlation between features.

### Tumor immunogenicity, cytolytic immune activity, and patient survival

Consistent with previous reports of the strong predictive power of immune cytolytic activity in checkpoint treated patients^2,8^, we found that the gene expression signature of CYT was significantly associated with survival in the non-checkpoint treated patients in SKCM and NSCLC (**Fig. 3b,d**). In order to assess if tumor fitness predicted survival in these cohorts, we computed maximal clonal tumor fitness scores in each patient (using the same method, software versions, and parameter specifications as *Luksza et al.*; see Methods) and found that they did not separate overall survival in either SKCM or NSCLC cohorts (**Fig. 3a,c**). As observed in other studies^9^, it is challenging for neoepitope based techniques to predict patient survival better than TMB alone, particularly when regressing out powerful covariates such as clinical tumor stage. To establish the relative predictive power of tumor fitness, we constructed a series of multivariable Cox proportional hazard models for each cohort which included TMB, tumor stage, Ϊ, and CYT. For each model, we computed the time dependent Brier score (see Methods; briefly, the weighted average of the squared distances between the observed survival status and the predicted survival probability of a model) and compared the resulting prediction error curves. We compared the relative predictive power of different regression models via bootstrap resampling. As shown in Figure 3, tumor fitness does not significantly reduce the prediction error in NSCLC nor SKCM compared to models with TMB alone, while CYT does so in SKCM but not in NSCLC (**Fig. 3e,f**). Computing the integrated Brier scores under cross-validation confirms these results are robust.

**Figure 3:**
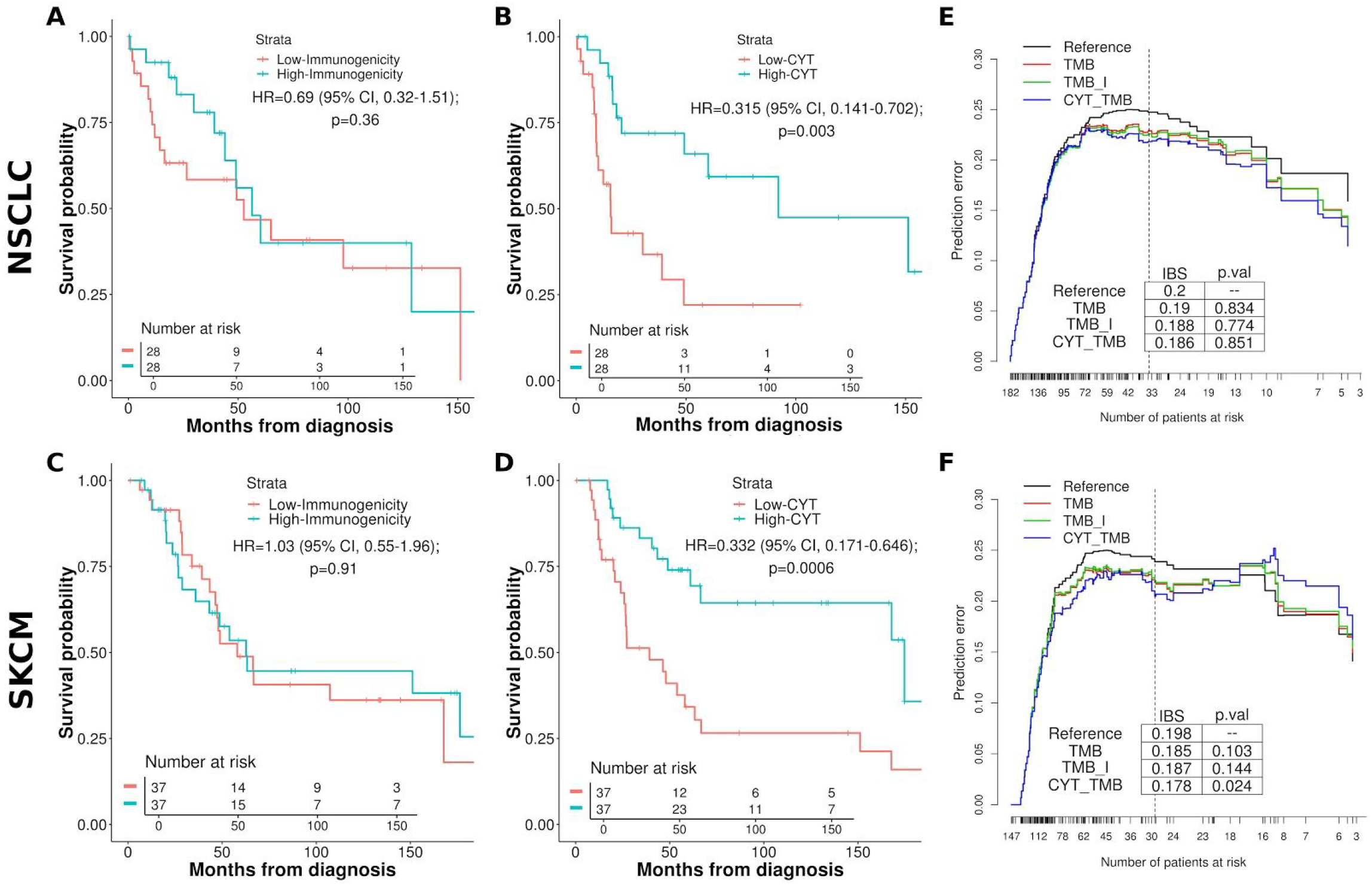
Univariable KM survival curves between patients grouped by highest and lowest immunogenicity (Ϊ) and immune activity marker expression (*CYT*) scores for the **A,B)** lung (NSCLC) and **C,D)** melanoma (SKCM) TCGA cohorts. Quantiles of *I* and *CYT* patient scores used to group cohorts with the lowest and highest groups represented by the curves; **E,F)** Time dependent Brier scores (prediction error curves) comparing multivariable cox proportional hazard survival models for the NSCLC cohort (E), and SKCM cohort (F). Reference is intercept, all other models include tumor stage covariates. TMB includes patient tumor mutation burden, TMB_I includes immunogenicity and TMB, CYT_TMB includes CYT and TMB. Vertical lines demarcate approximately 80% of the patients, for which integrated Brier scores were calculated.

### Neoepitopes, not viral epitopes, better bind MHC-I and also associate with immune response in HBV-related liver cancer

To examine the relationship between TCR recognition and neoepitope sequence similarity to pathogen derived epitopes in samples that coexpress both, we examined patients with hepatocellular carcinoma co-infected with HBV from the LIHC TCGA dataset. We assessed 190 patients with matched WES and RNA-Seq data, detecting HBV transcripts in 115/190 (60%) via HBV genome assembly from RNAseq data unmapped to human genes (see Methods). Of these, 88 patients produced HBV matching protein sequences through ORFfinder. Using RNA-seq data for patient HLA typing, we estimated the distribution of binding affinities of the viral epitopes and compared it with that of neoepitopes produced by tumor mutations. A total of 133,834 unique tumor neoepitopes and 30,357 HBV viral epitopes were predicted across the patients. Cumulative distribution functions of the predicted MHC-I binding affinities indicate a significant MHC-I binding affinity preference for tumor derived neoepitopes over that of HBV viral cofactors (**Fig. 4**). This result is robust against subsampling among neoepitopes (**Supp. Fig. 1**). In addition, we found that tumor neoepitope burden is marginally but significantly associated with patient TIL burden (Spearman’s ρ=0.24, p=0.038), but HBV epitope burden was not (restricting to MHC-I strong binders, ic50 < 200 nM; **Supp Fig. 2**), further suggesting that TIL recruitment is preferentially driven by relative neoepitope burden compared to viral burden. Our observation forms a counterexample to the model assumption that viral sequence similarity drives preferential epitope recognition.

**Figure 4:**
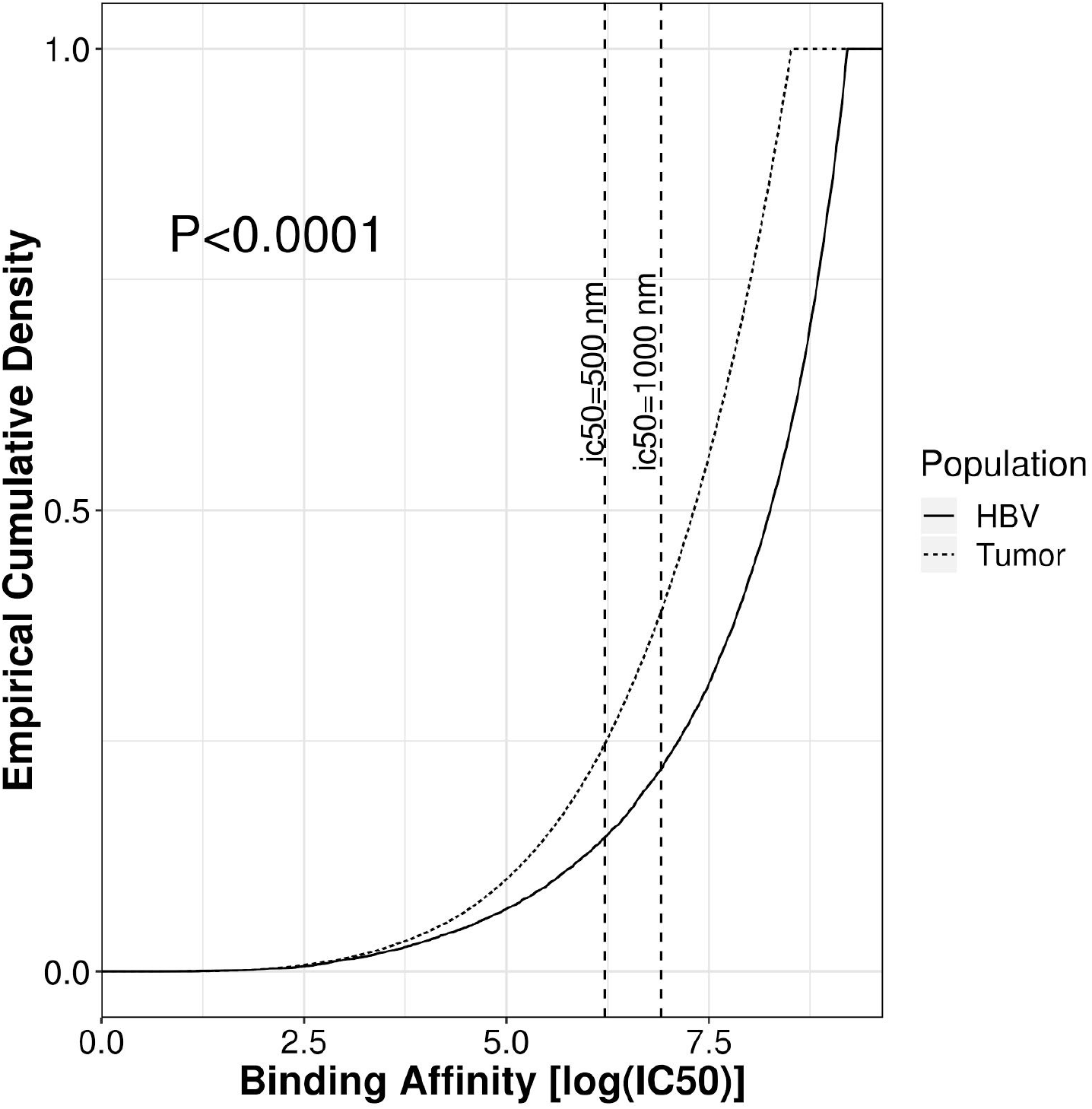
Distribution of MHC-I binding affinities for tumor neoepitopes (dotted line) and HBV epitopes (solid line). Smaller ic50 values indicate a stronger binding affinity. Tumor neoepitopes are on average more likely to bind to patient MHC-I alleles than epitopes from HBV proteins.

## Discussion

Determining the degree of ‘non-selfness’ of mutant peptides in the face of enormous individual variability of TCR repertoires is notoriously difficult, and yet critical, to answer at scale. Point somatic mutations located outside the MHC-binding anchor positions may appear to be more ‘non-self’ since they may lead to significantly increased peptide-MHC binding compared with wildtype sequence. However it has traditionally not been straightforward to translate this notion into a better survival or treatment response predictor. The relative spectra of subclonal and clonal immunogenic mutations, and similarly mutant sequence homology with experimentally validated pathogen epitopes, potentially contain important tumor evolution signatures which may indicate intriguing similarities with pathogen evolution.

To gain further intuition into a recent neoepitope based tumor fitness scoring model^4^, we applied it to samples from checkpoint-treatment-naive samples of the same tumor types in the TCGA. A key aim was to test how this model predicts survival given that it has shown promise in predicting response to immune checkpoint inhibitors. Although this setting may involve substantially different (thus far uncharacterized) tumor-immune dynamics, we reasoned that a sufficiently general notion of tumor fitness should correlate with patient survival, especially in those cohorts where TIL and CYT are already known as strong survival correlates.

Our results suggest that the proposed tumor fitness measure based on pathogen epitope similarity does not correlate with CYT in either TCGA NSCLC nor SKCM, potentially indicating that it is specific to the patients treated with checkpoint inhibitors. Furthermore, in a test of a key model assumption, in the case of HBV-related HCC in the TCCA LIHC, we find that even when viral epitopes are competing with neoepitopes for T cell presentation there is actually a substantial average bias towards higher MHC-I binding affinity for the neoepitopes. In addition, the significant correlation between TMB and TIL burden further suggests that the tumor dominates immune recruitment over HBV viral epitopes. It is interesting to speculate on whether other tumors with known viral cofactors, such as the Human Papillomavirus in cervical or head and neck cancers, and Epstein Barr virus in Burkitt’s lymphoma, show similar biases.

Furthermore, we find that tumor fitness in the treatment-naive samples does not improve prediction error of survival beyond that of TMB and clinical tumor staging alone, while a simple measure of CYT does so in SKCM. The tumor fitness model has two key degrees of freedom, ɑ and k, that were tuned to reduce prediction error in patients treated with checkpoint inhibitors^4^. However, given that the alignment score distributions for the TCGA cohorts were not significantly different from the matched checkpoint treated settings (**Supp. Fig. 4**), we used the same values in our modeling of these treatment-naive samples.. While these data suggest that the tumor immune dynamics in treated patients are notably different than in untreated patients, it may also simply reflect a strong dataset-specific dependence of the tumor fitness score. Indeed, as noted the CYT measure alone seems to reduce survival prediction error, as found by others^10,11^ and *Luksza et al.* themselves as a useful component to add to the tumor fitness score. Though the improvement of survival prediction by the inclusion of CYT may be, essentially, expected based on its efficacy as an independent predictor of therapy responsiveness in the checkpoint-setting^8^, we demonstrate here that this common-sense expectation holds true in the non-checkpoint setting as well.

Although our study points out key limitations in extending the paradigm of tumor fitness scoring to untreated patients, it does have the same caveats any in-silico immunogenicity estimation based analysis method has. Indeed, the accuracy of in-silico predictions of neoepitopes or viral binding affinity in estimating immune reactivity are at best suboptimal compared to using mass spectroscopy. Also, in comparing viral and neoepitopes it’s important to note that several lines of evidence suggest that the replication status of HBV is dependent on the differentiation status of the cells^12,13^. Thus, the virus may hardly replicate in HCC cells and further reduce the viral epitope burden within the tumor.

Overall our study indicates that while tumor fitness scores can be used to predict response to checkpoint inhibitors, they do not seem to predict survival in untreated patients where endogenous immune activation is a significant survival factor. It is important to note that while this tumor fitness measure and its underlying assumptions have been adapted with positive predictive results in other contexts^14,15^, we find that the exact form of the model is not predictive in these endogenous cases. Nevertheless, it is encouraging to observe that CYT measures are powerful in both treated and untreated patients, which may point to the existence of universally predictive and prognostic immune activation signatures.

## Methods

### TCGA analysis

RNA-Seq data for the TCGA patient cohorts was aligned to Hg38 with STAR (v2.5.1b) in two-pass mode^16^. Gene counts for Gencode v23 (www.gencodegenes.org) gene annotations were generated using featureCounts. Read counts underwent TMM normalization and logCPM transformation using voom^17^. Mapped reads were used to allelotype (MHC class-I loci) each patient^18^, and estimate the putative TIL burden and clonality per patient by profiling TCR and BCR sequences with MiXCR^19^, normalizing by patient library size. To predict neoepitopes and associated viral epitope burden, we used Topiary^20^ to call mutation-derived cancer T-cell epitopes from somatic variants, tumor RNA expression data, and patient class I HLA type. This tool matches mutations with gene annotations, filters out non-protein coding changes, and finally creates a window around amino acid changes, which is then fed into NetMHCCons for each patient HLA allele across tiles of 9-12 aa in length. Given that HLA-I processes neoepitopes by degradation to non-conformational 8-11 amino-acid residues, we excluded neoepitopes with mutations obscured to T-cells within HLA-I binding pockets (anchor residues at 2,9 aa that are mutated with respect to the wild-type form of the epitope). In the case of frameshift mutations, in principle this window starts from the mutation minus the length of the peptide up to the first stop codon. Neoepitopes were filtered for noise using a binding affinity threshold of ic50 < 10,000 nM, which removed 89% of predictions. A measure of tumor mutation burden (TMB) for each patient was calculated by enumerating the number of called DNA-mutations in patient associated MAF files.

### Tumor fitness score

We assigned tumor immunogenicity scores following the method outlined by *Luksza et al.* Briefly, a protein BLAST of the neoepitopes from the SKCM and NSCLC cohorts was performed per patient against a database of viral epitopes sourced from the IEDB^5^; the best hits with higher sequence similarity normalized by sequence database size (bitscore) for each neoepitope were retained. A second, randomized set of neoepitopes was also blasted, as a negative control group. Smith-Waterman alignments scores (|s,e|) were calculated for each match, following a strict 500 nM binding affinity threshold filtering for all predicted neoepitopes. For each patient, a TCR recognition potential (*R*) was calculated by modeling the probability for a given neoepitope to be bound to a viral match from the IEDB using:

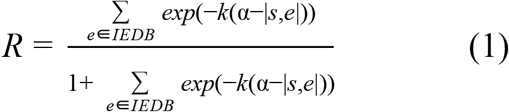

with constants ɑ (binding curve displacement) and k (curve steepness) retained from the original model in matching to each TCGA patient cohort, after comparisons of alignment score distributions with checkpoint treated cohorts revealed no significant differences (**Supp. Fig. 4**). For each neoepitope, we calculated an MHC-I binding affinity amplitude score (*A*), as a ratio of the binding affinities of the mutant neoepitope and the wild type epitope, which were corrected for dissociation constant bias by multiplying by the reciprocal max among all wild type epitopes. Tumor immunogenicity was assigned by selecting the maximum product of these scores from among all neoepitopes across patient clones, *j*:

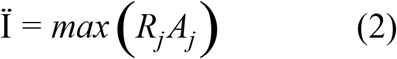

This term represents the effective expected size of immune response and is used to rank the dominant immunogenic clone per patient. The inverse of this score, tumor fitness, describes the likelihood of evading immune detection.

Cytolytic T-Cell activity (CYT) scores were calculated using the geometric mean of the RPKM expression of granzyme A (*GZMA*) and perforin-1 (*PRF1*)^6^.

### HBV antigen calling

Raw RNA sequencing reads that did not map to the GRCh38 reference genome was assembled into contigs using Trinity (using *–no_run_chrysalis –no_run_butterfly* flags, which only invokes the Inchworm step) to perform greedy kmer-25 contig assembly^21^. Resulting contigs with a sufficiently high entropy (to exclude homopolymer sequences), at least 100 bp long and supported by at least 20 reads were retained and aligned with BLAST against the HBVdb^22^ reference genomes (e-value < 1e-6) for HBV (genotypes A, B, C, D, E, F, and G), effectively identifying the HBV positive patients (n=115). CD-Hit was used to consolidate highly similar contig sequences (e-values < 1e-10)^23,24^. Finally, ORFfinder was used to predict the putative HBV protein products from each contig, which were validated and filtered (e-value < 1e-10) by a protein BLAST against an HBV protein database compiled from HBVdb for the HBV genotypes above^25^. Importantly, our contig length filtering means that ORFs shorter than 300 aa are excised. HBV epitopes were predicted using the matching protein products and patient HLA-typing calls with Topiary (see above), and predicted peptides were filtered for noise by binding affinity (ic50 < 10,000 nM), reducing predictions by 94%.

### Survival Analysis

Kaplan-Meier survival curves and risk tables using patient tumor immunogenicity, Ϊ, and immune activity, CYT, were created with the survival and survminer packages^26,27^. Patient grouping into low and high tumor immunogenicity and immune activity was done using the bottom and top 15% quantiles for these measures, respectively. Cox proportional hazard models using tumor stage and TMB, Ϊ, and CYT covariates were constructed using the rms package^28^.

### Statistical Analysis

Significance for all test measures were determined at p-values < 0.05. Spearman correlations between expression based tumor immune measures (CYT, TIL burden, TIL clonality) and TMB and immunogenicity score (Ϊ) were calculated to correct for outliers. We used a one-sided Kolmogorov-Smirnov test compare viral and neoepitope MHC-I binding affinity distributions, with an alternative hypothesis that the neoepitope distribution falls above the viral distribution. Additionally, to assure that the MHC-I binding affinity bias toward tumor neoepitopes was not confounded by a larger epitope set, we subsampled 10,000 epitopes from both populations and compared binding affinity distributions by one-sided (neoepitope greater) Kolmogorov-Smirnov test for 1,000 replicates (**Supp. Fig. 1**).

To address possible violation of the assumption of proportional hazard ratios between populations in our survival models, we computed the correlation between survival covariates (Ϊ, CYT) and time for each model. We found all model covariates were time-independent, and thus a log-rank test was used to evaluate significance in predicted survival outcomes between groups for Kaplan-Meier survival estimates. The relative predictive power of various Cox survival models was evaluated using the pec R package^29^ by computing the Integrated Brier Score (IBS) across .632+ bootstrap resampling with 100 iterations (*B*=100). We used the Kaplan-Meier estimator for the censoring times in the bootstrap simulations. The weights in the IBS correspond to the probability of not being censored. Model error curves were compared using Wilcoxon test for IBS comprising approximately 80% of total patients, with test significance provided in graphic tables (**Figure 3e,f**). Error prediction reduction measured by IBS with respect to time from diagnosis (5 years) was also evaluated (**Supp. Fig. 5**). IBS cross validation was performed by sampling Brier scores with 100 iterations across random patient groups for each model and applying a one-sided (*TMB_I* less) Kolmogorov-Smirnov test.

## Availability of Code and Data

All code and non-protected data is available upon reasonable request to the corresponding author.

## Author Contributions

B.L. conceived and directed research. N.A. and A.B. performed principal data analysis. A.B., A.V., E.G.K. and B.L. wrote and edited the manuscript.

## Additional Information

The authors declare no competing interests.

