## Supplemental Materials for "Tumor fitness, immune exhaustion and clinical outcomes: impact of immune checkpoint inhibitors"

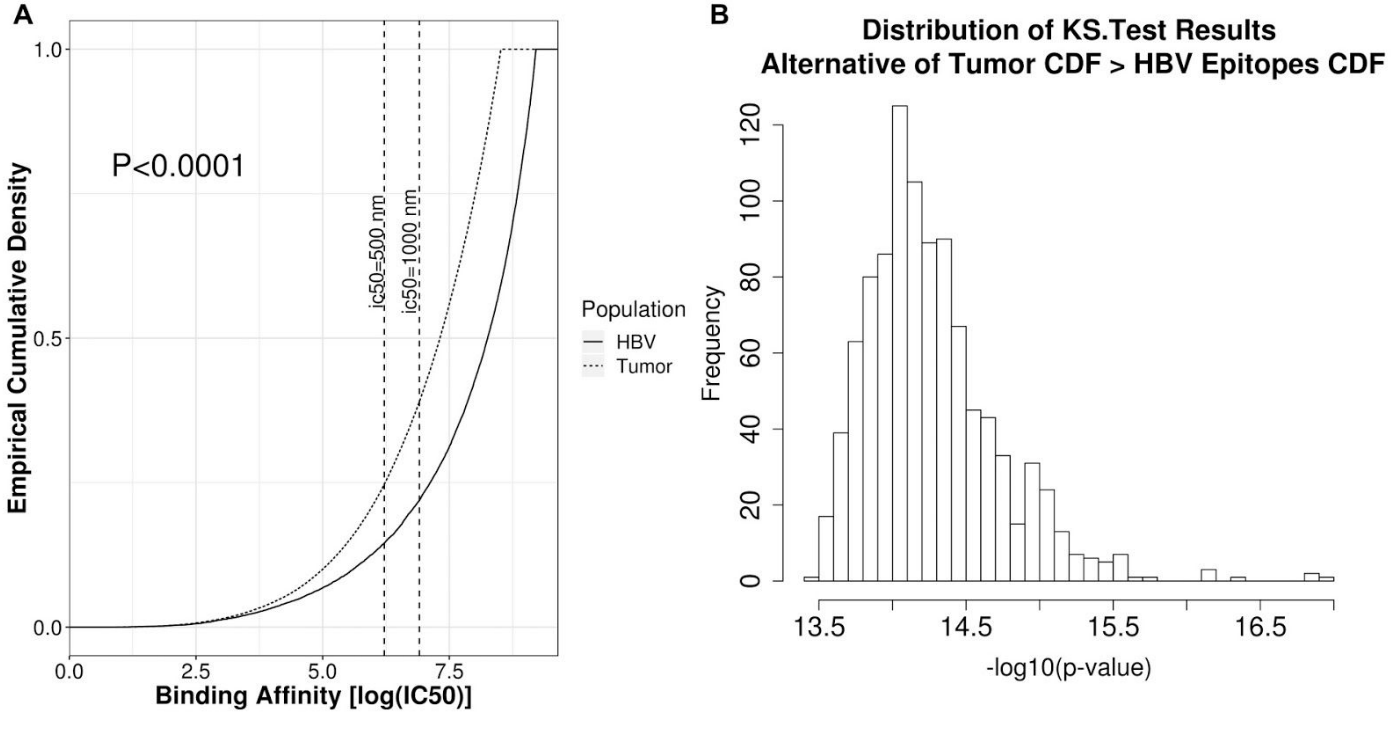


**Supp. Figure 1:** Kolmogorov-Smirnov test results (**A**) of randomly subsampled distributions of tumor neoantigen binding affinities compared to HBV antigens across 1,000 iterations. **B)** Distribution of test p-values strongly support the alternative hypothesis that tumor neoantigen cumulative distributions were significantly less than HBV distributions, indicating better average binding affinity scores are robust to subsampling.

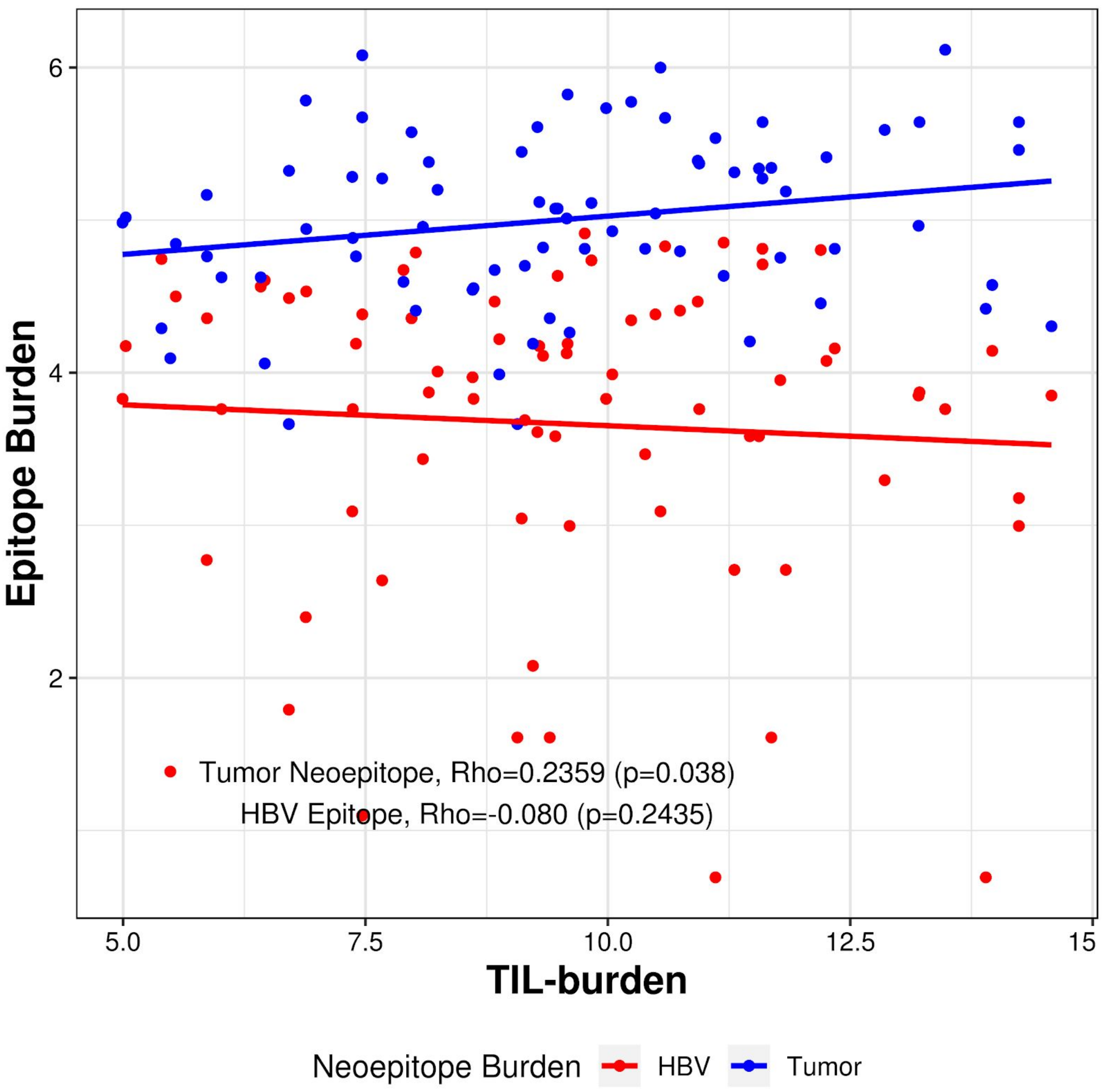


**Supp. Figure 2:** Tumor neoantigen burden is slightly but significantly positively correlated with patient TIL-burden, while HBV antigen load is not. Spearman correlations are presented for epitope group and TIL-burden associations.

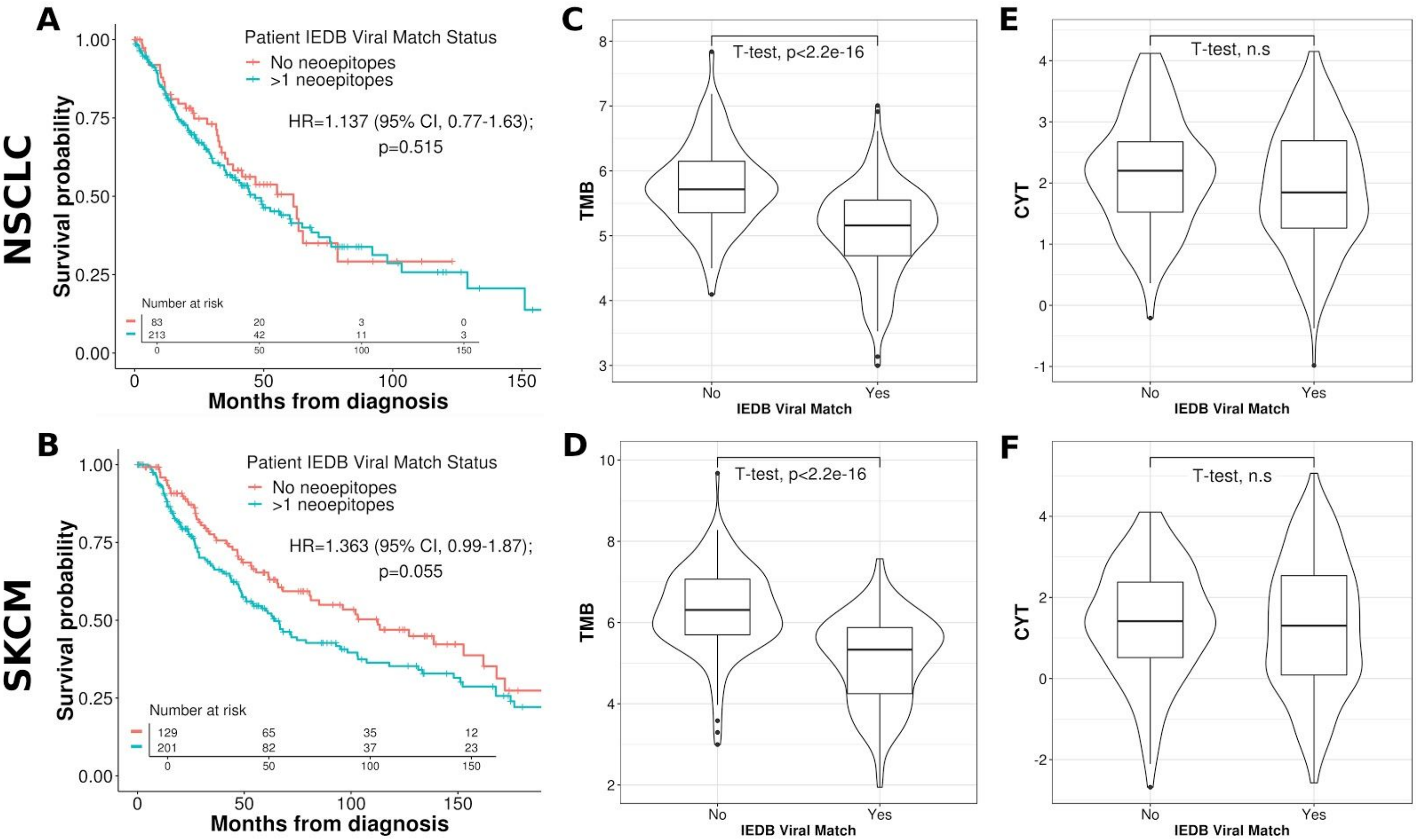


**Supp. Figure 3:** Kaplan-Meier survival curves between patients with and without neoepitope BLAST matches against the viral IEDB database for the **A)** lung and **B)** melanoma TCGA cohorts. Tumor mutational burdens for patients with viral neoantigen matches fall significantly below those with no matches in both the **C)** lung and, **D)** melanoma patient groups (two-tailed T-test). Differences in cytolytic activity (CYT) expression were found to be negligible for both the NSCLC and SCKM patients between those with and without IEDB matches (**E** and **F**, respectively) (two-tailed T-test).

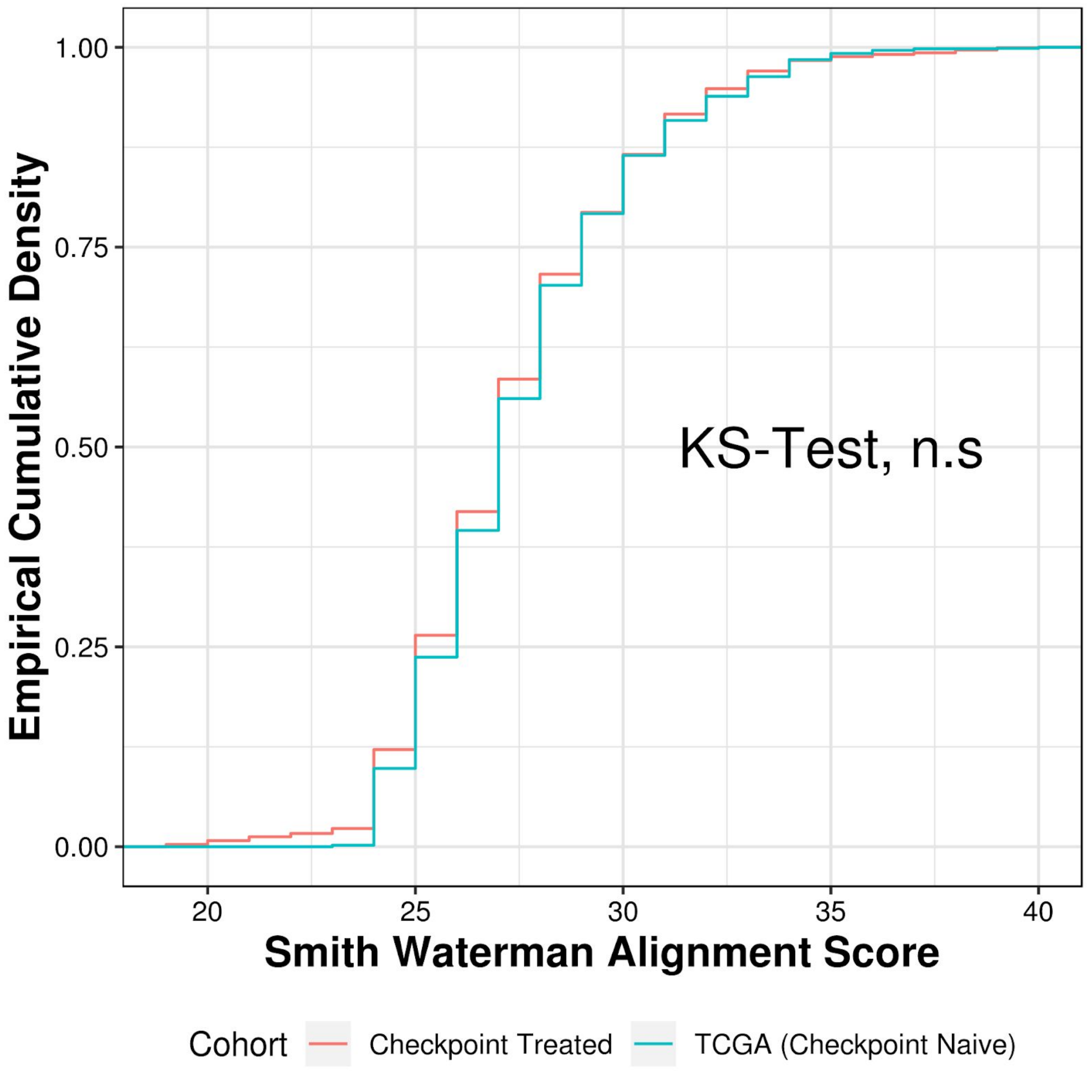


**Supp. Figure 4**: Comparison of smith-waterman alignment score distributions for tumor neoepitopes to best matching viral epitopes from the TCGA lung and melanoma cohorts and the checkpoint treated patient cohorts used in *Luksza et al*. By Kolmogorov-Smirnov two-sided test, no significant difference between alignment score distributions was observed.

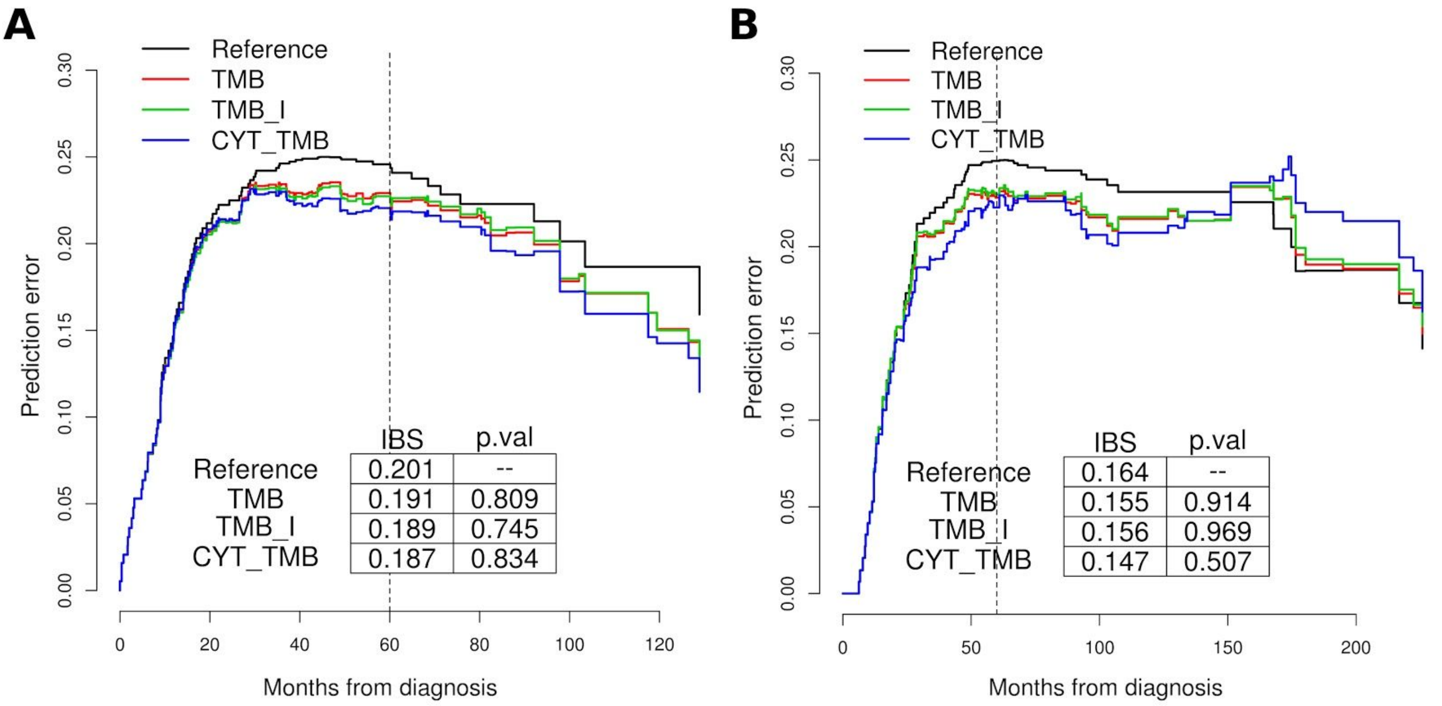


**Supp. Figure 5**: Prediction error curves for Cox survival models in **A**) lung, **B**) melanoma TCGA cohorts using covariates TMB, TMB and Immunogenicity (I), and TMB and CYT, with respect to months from patient diagnosis. Time dependent integrated Brier scores were evaluated at 5 year survival from diagnosis (60 months). Under this evaluation criteria, no models significantly reduced prediction error over the naive reference model.
